# EphA2 and Ephrin-A1 Utilize the Same Interface for Both *in cis* and *in trans* Interactions That Differentially Regulate Cell Signaling and Function

**DOI:** 10.1101/2025.07.31.667925

**Authors:** Soyeon Kim, Paul Toth, Wei Wang, Elena Seiradake, Xiaojun Shi, Bingcheng Wang

## Abstract

The 14 members of Eph receptor tyrosine kinases (RTK) bind to membrane-tethered ligand called ephrins and mediate cell contact signaling where the receptors and ligands engage *in trans* on adjacent cells. Previous studies reveal that some Eph and ephrin pairs are coexpressed on the same cells, including EphA3-ephrin-A3 and EphA4/ephrin-A5, can also interact with each other *in cis*. However, significant discrepancies persist as to the molecular basis and functional significance of the *cis* interactions, owning to the difficulties to directly interrogate the interactions. Here, we utilize time-resolved live cell fluorescence spectroscopy to demonstrate direct *in cis* interactions between EphA2 and ephrin-A1. Structure-guided mutagenesis mapped interactions to two salt bridges between the ligand- and receptor-binding domains of EphA2 and ephrin-A1. Interestingly, the same interface is shared with *in trans* interaction. Consequently, EphA2-ephrin-A1 interaction *in cis* competes with their interaction *in trans*, which leads to attenuation of EphA2 canonical signaling and inhibition of cell rounding when ligand is presented *in trans*. EphA2 and ephrin-A1 are widely coexpressed in many epithelial tissues, and dysregulation of their expression is known to contribute to tumor initiation and progression. The detailed molecular characterization of the mutually exclusive *cis* and *trans* interactions uncovers a new mechanism underpinning their unique roles in oncogenesis.

**Significance Statement:** EphA2 exerts dual functions in tumorigenesis, depending on the binding status of its membrane-tethered ephrin-A ligands. Ligands presented *in trans* on adjacent cells activate EphA2 canonical signaling associated with tumor suppression, whereas loss of ligand expression promotes oncogenic noncanonical signaling of EphA2 via serine 897 phosphorylation. Combining time-resolved spectroscopy in live cells, structure-guided mutagenesis, we show strong in *cis* interactions between EphA2 and ephrin-A1, which shares the same interface as interaction in trans. Moreover, the *cis* interaction interferes with ligand binding *in trans*, attenuates EphA2 canonical signaling. Our results uncover a new mechanism of EphA2 regulation by its co-expressed ligand ephrin-A1 with important implications in its known roles in oncogenesis as well as other disease processes including development of cataract.

## Introduction

The 14 members of Eph receptor tyrosine kinases (RTK) and their membrane-tethered called ephrins engage *in trans* on surface of adjacent cells to mediate juxtacrine signaling that regulates diverse developmental and pathological processes including Alzheimer disease and cancer (1, 2). Extensive in vitro and in vivo studies show that there is a significant overlap of expression with each cell type possessing distinct repertoire of ligands and receptors, leading to the long-standing question whether and how they interact on the membrane of the same cells, and more importantly, the functional significance of such *in cis* interactions. Supporting this notion, multiple reports suggest that Ephrin ligands can also interact with Eph receptors *in cis,* when they are co-expressed on the same cells (3–8). However, the molecular details of these interactions remain to be fully elucidated due to discrepancies in literature and technical difficulties examining such interactions under physiological conditions. Several reports suggest that *in cis* interaction is mediated by ephrin-A3 binding to the FN1-FN2 domains of EphA3 and EphA4 (3–8). However, a later study argues that it may be mediated by the ligand-binding domain (LBD) of EphA4 (9). The divergent conclusions are drawn from receptor truncation and antibody blocking approaches, leaving the molecular details for the *in cis* interactions yet to be fully elucidated. Further complicating the debate, some researchers have observed spatial segregation of co-expressed Eph and ephrins into distinct membrane microdomains, raising doubts about the very existence of *cis* interaction (11). Crystal structure suggests ephrin-A5 interaction with EphA2 FN2 domain (10), although the actual involvement of putative tripartite interface in the EphA2-eprhin-A5 interaction has not been experimentally examined.

EphA2 has been extensively investigated in cancer due to its overexpression in late-stage human solid tumors that are associated with poor prognosis. Cellular, biochemical, and genetic studies reveal that EphA2 can function as either a tumor suppressor or an oncogenic protein, depending on the ligand-binding status. When stimulated with its ligand, the catalytically activated EphA2 is autophosphorylated on tyrosine residues, triggering canonical tumor suppressive signaling including the inhibition of Ras/MAPK (12), PI3K/Akt, and integrin inside-out signaling pathways (13). Conversely, in the absence of ligand, EphA2 becomes phosphorylated at serine 897 by AGC family serin/threonine kinases, which transform EphA2 into an oncogenic protein that drives tumor invasion and metastasis (1, 2, 14, 15).

Our recent study (16) elucidates the molecular mechanisms governing EphA2 self-assembly and activation. We used a technology called the Pulsed Interleaved Excitation Fluorescence Cross-Correlation Spectroscopy (PIE-FCCS) that enables time-resolved live cell measurements of protein-protein interactions. We found that the ligand-free EphA2 is auto-assembled in multimers. Three interfaces in the ectodomain mediate the multimeric assembly, ligand-binding domain (LBD) to LBD, Sushi domain to Sushi domain (collectively referred to as Head-to-Head interfaces), and FN2 domain to LBD domain (Head-to-Tail interface). In most normal epithelia, EphA2 is co-expressed with ephrin-A1, a major cognate ligand that has been extensively characterized for regulating EphA2 signaling and function in vitro and in vivo (2, 16, 17) . As such, the role of the *in cis* interaction is particularly important in understanding EphA2-ephrin-A1 signaling in epithelial homeostasis and malignant transformation, which has not been closely examined and molecularly characterized.

In this study, we addressed these inconsistencies using a fluorescence-based biophysical method capable of observing interactions between proteins of interest in live cells. Our results demonstrate that EphA2 and ephrin-A1 interact in *cis* mainly through the LBD-RBD interface. We also show that this *in cis* interaction competes with *in trans* interactions, thereby inhibiting the tumor-suppressive functions of EphA2.

## Results

### Live cell fluorescence correlation spectroscopy revealed EphA2 interaction with ephrin-A1 *in cis*

Pulsed Interleaved Excitation Fluorescence Cross-Correlation Spectroscopy (PIE-FCCS) is a powerful platform that can measure the mobility of membrane proteins in live cell membranes and probe their interactions (**Fig. 1A**). Details of this methodology have been reported previously (16). Briefly, two spatially overlapping and time-interleaved laser beams are focused on the small area of the peripheral cell membrane to excite GFP- and mCherry-tagged proteins co-expressed in the same cell (**Fig. S1A**). The emitted photons are collected and mathematically transformed into autocorrelation or cross correlation curves (**Fig. S1B,C**). Two parameters can be derived from the correlation curves, fraction correlated (*f_c_*) that reflects whether the membrane proteins are interacting or not, and diffusion coefficient (D) that correlates with the sizes of the complex (13) (**Fig. S1D**). Zero or near-zero *f_c_* indicates that there is no interaction between the proteins (**Fig. S1B**). Positive *f_c_*values higher than 0.1 indicate that the proteins interact (**Fig. S1C**). To investigate if EphA2 and ephrin-A1 may directly interact in live cells using PIE-FCCS, we coexpressed the two proteins that were differentially tagged with GFP or mCherry (mCh) in Cos-7 cells (**Fig. 1B**). As expected, GFP-EphA2 and mCherry-ephrin-A1 colocalized at the cell-cell contact sites through interactions *in trans* (**Fig. 1B**, while boxes). To ensure only *cis* EphA2-ephrin-A1 interaction is measured by PIE-FCCS, the lasers are focused only on areas (**Fig. 1B**, white arrow) that are away from cell-cell junctions.

**Figure 1.**
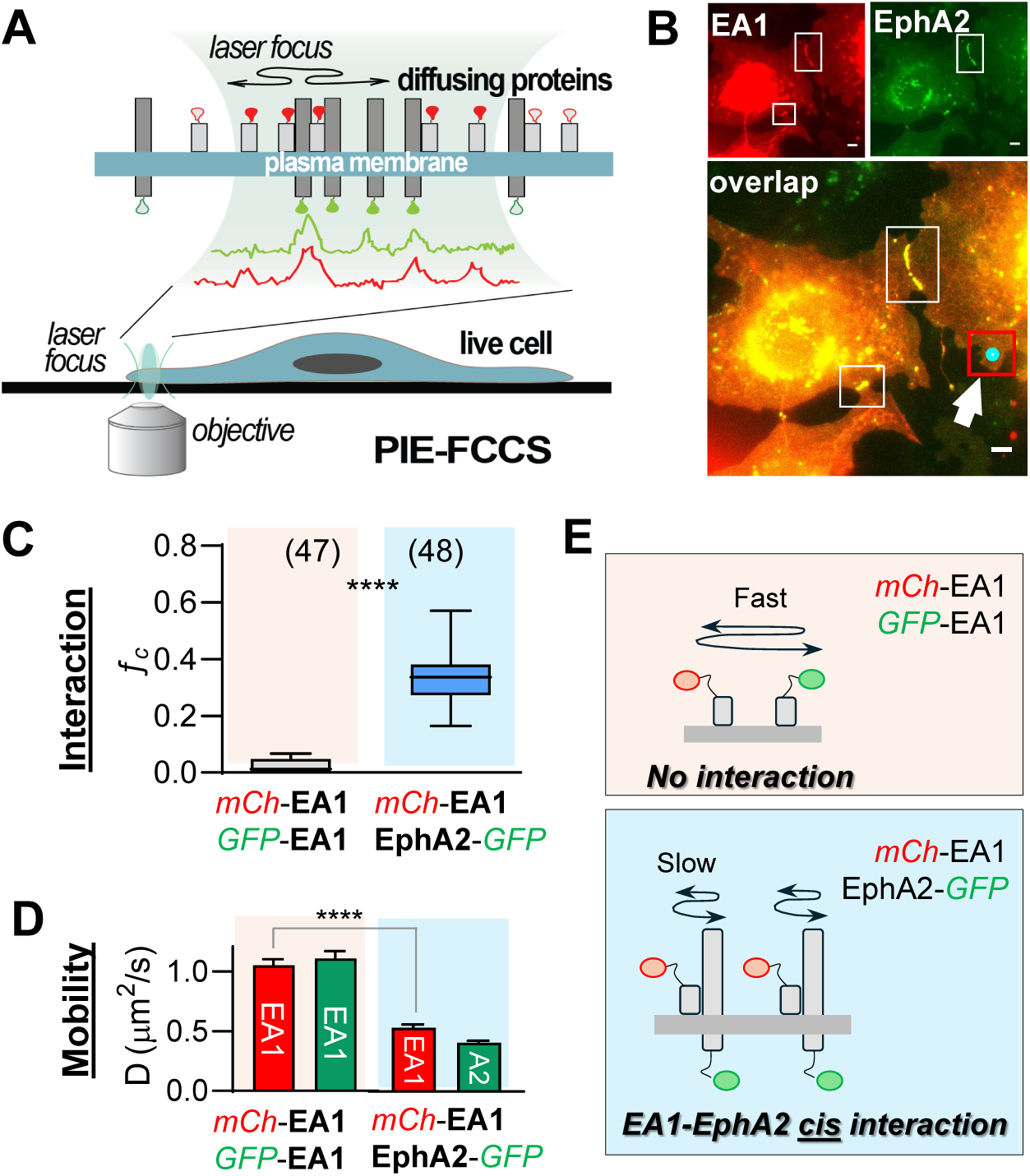
A) Schematic depiction of experimental setup of PIE-FCCS. B) Sample images of cells co-expressing mCh-EA1 and EphA2-GFP. White boxes highlight the cell junction where EA1-EphA2 trans-interaction is, while white arrow point to where the laser focus (indicated with cyan dot in the red box) is to ensure only cis-interaction is detected. C) Left: fc values and D) diffusion coefficients of mCh-EA1 co-expressed with either GFP-EA1 (pink area) or EphA2-GFP (blue area). E) Schematic depiction of monomeric EA1 with fast diffusion (top) and slower diffusion of EA1 undergoing cis interaction with EphA2 (bottom). The box plot represent third quartile, median, and first quartile, and the whiskers indicate the 10th to 90th percentile. The total cell number that was used for each sample is reported at the top of the box plots. Data were analyzed by T-test; ****p < 0.0001, **p < 0.001, *p < 0.01.

In keeping with our previous report (16), there was no homotypic interaction between mCh-ephrin-A1 and GFP-ephrin-A1 coexpressed in the same cells, as reflected by the near-zero *f_c_* values (**Fig. 1C**, pink shade), suggesting that ephrin-A1 exists as a monomer in live cell membranes. By contrast, when mCh-ephrin-A1 was co-expressed with EphA2-GFP, high *f_c_* values were recorded (**Fig. 1C**, blue shade), indicating strong EphA2-ephrin-A1 interactions. This is further supported by the slower mobility of EphA2-ephrin-A1 complex compared to the high mobility of monomeric ephrin-A1 (**Fig. 1D**). Neither mCh-ephrin-A1 nor EphA2 GFP showed molecular density-dependent change in *f_c_* values (**Fig. S1E, F**). Diagrams depicting the monomeric ephrin-A1 and EphA2-ephrin-A1 complexes are shown in **Fig. 1E**.

### EphA2 and ephrin-A1 *cis* interaction is mediated by two salt bridges between receptor binding domain (RBD) of ephrin-A1 and ligand-binding domain (LBD) of EphA2

Our previous report using X-ray crystallography demonstrated that EphA2 ligand-binding domain (LBD) and ephrin-A1 interact through a “lock and key” mechanism *in trans* (18). The rigid D-E and J-K loops of EphA2 and G-H loops of ephrin-A1 interlock, stabilized by two salt bridges. We posited that the same interface mediating the strong receptor-ligand interaction *in trans* could also be involved *in cis* interaction between ephrin-A1 and EphA2 co-expressed on the same cells. To test this, we created the K107E and E119K mutations on ephrin-A1 (KE/EK) to disrupt the salt bridges with D53 and R103 of EphA2 (**Fig 2A**). We first verified if the mutation impaired the ephrin-A1 binding to EphA2 *in trans*. To this end, we expressed mCherry-tagged ephrin-A1 WT (mCh-ephrin-A1) or mutant (mCh-ephrin-A1^KE/EK^) in HEK293 cells, and cocultured them with HEK293 cells expressing EphA2-GFP, and analyzed EphA2 activation via western blot (19)(**Fig. 2B**). Coculture of EphA2-expressing cells with ephrin-A1^WT^-expressing cells was expected to enable receptor-ligand interactions *in trans* upon cell-cell contact. Indeed, we observed strong EphA2 activation as indicated by the high pY-EphA2/total EphA2 ratio (19), which was accompanied by dramatic degradation of the receptor (**Fig. 2B**). By contrast, mCh-ephrin-A1^KE/EK^ was unable to cause EphA2 activation and degradation, confirming the mutant ephrin-A1 effectively disrupted *in trans* interactions. Catalytically activated EphA2 is known to undergo rapid endocytosis that is observed in cells expressing WT-but not the ephrin-A1^KE/EK^ (**Fig. S2**).

**Figure 2.**
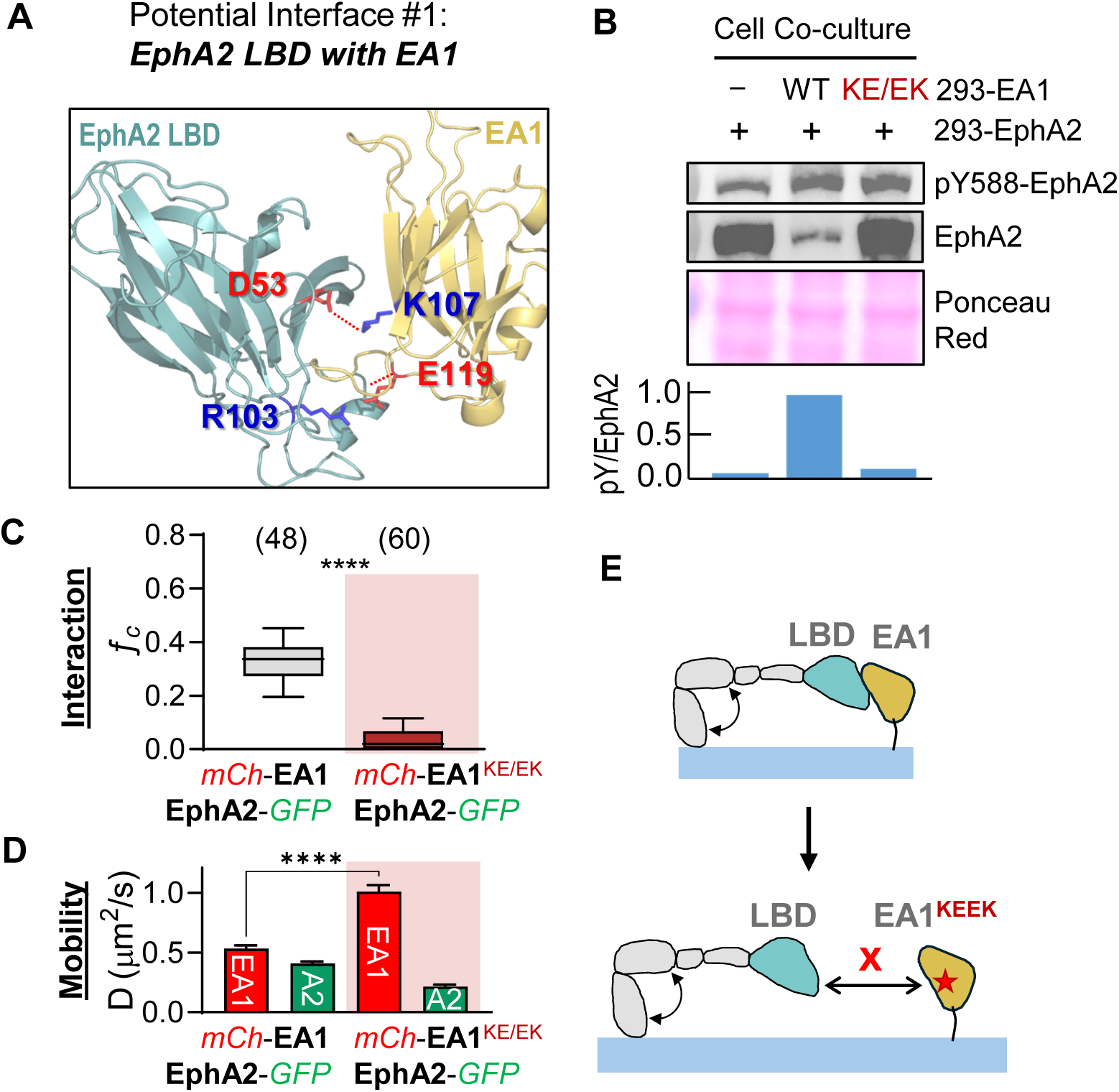
A) Structural depiction of EphA2-Ephrin-A1 cis interaction via EphA2 LBD. Two pairs of salt bridges are highlighted. B) Western blots showing that EA1KE/EK, which harbors mutations to disable the binding with LBD, cannot activate EphA2. C,D) fc values (C) and diffusion coefficients (D) of EphA2-GFP co-expressed with either mCh-EA1 (white area) or mCh-EA1KEEK (red area). EA1KEEK abolishes the cis interaction with EphA2 entirely, as depicted in the cartoon D. E) Cartoon depiction of EA1-EphA2 cis interaction via EphA2 LBD. The cis interaction is disrupted by the KEEK mutation. The box plots in panel C represent third quartile, median, and first quartile, and the whiskers indicate the 10th to 90th percentile. The total cell number that was used for each sample is reported at the top of the box plots. Data were analyzed by T-test; ****p < 0.0001, **p < 0.001, *p < 0.01.

Next, we coexpressed mCh-ephrin-A1^KE/EK^ with EphA2-GFP and examined the *cis* interaction using PIE-FCCS on sparsely plated cells to minimize cell-contact-induced interaction *in trans*. Compared with ephrin-A1^WT^ that interacted with EphA2 displaying an *f_c_* value 0.32, ephrin-A1^KE/EK^ mutant showed little *in cis* interaction, with the *f_c_* value reduced to near zero (0.02) (**Fig 2C**). The mobility of ephrin-A1^KE/EK^ increased compared with ephrin-A^WT^ (**Fig. 2D**, red), suggesting that ephrin-A1^KE/EK^ diffuse freely in the membrane untethered to the coexpressed EphA2. Unable to engage with ephrin-A1^KE/EK^ *in cis*, the co-expressed EphA2-GFP self-assembled into homotypic multimers as we reported previously (16), which likely accounts for the reduced EphA2 motility compared with cells expressing ephrin-A1^WT^ (**Fig. 2D**). Together, these results demonstrate that the *cis* and *trans* interaction between EphA2-ephrin-A1 utilize the same interface involving the two salt bridges as illustrated in **Fig. 2A**.

### The two type III fibronectin repeats of EphA2, FN1 and FN2, are not involved in interaction with ephrin-A1 *in cis*

The results above demonstrate that salt bridges between EphA2 LBD and ephrin-A1 RBD are the major drivers of the interactions *in cis*. However, the possibility of additional contributors cannot be fully ruled out. Indeed, FN1 and FN2 domains of EphA3 and EphA4 have both been previously shown to mediate *in cis* interactions with ephrin-A3 and ephrin-A2 (3–8). Supporting this notion, crystal structure of the EphA2-ephrin-A5 complex reveals a tripartite interface where ephrin-A5 RBD interacts with the FN1 and FN2 domains of EphA2 (10). The interface buries a large surface area and encompasses multiple contact points between EphA2 and ephrin-A5, which are proposed to mediate the interaction *in cis* (**Fig 3A**). Homology comparison and molecular modeling shows similar large contacts are also present in EphA2-ephrin-A1. To investigate whether the tripartite interactions may contribute to EphA2-ephriin-A1 interaction *in cis*, we created bulky glycosylation sites in the FN1 and FN2 to disrupt the interface. Since N-linked glycosylation occurs on asparagine in N-X-S or NXT motifs, T336N/V338T (N336) and V484N/V486T (N484) double mutations were generated to create glycosylation sites on N336 in FN1 and N484 in FN2, either individually or in combination (**Fig. 3A**).

**Figure 3.**
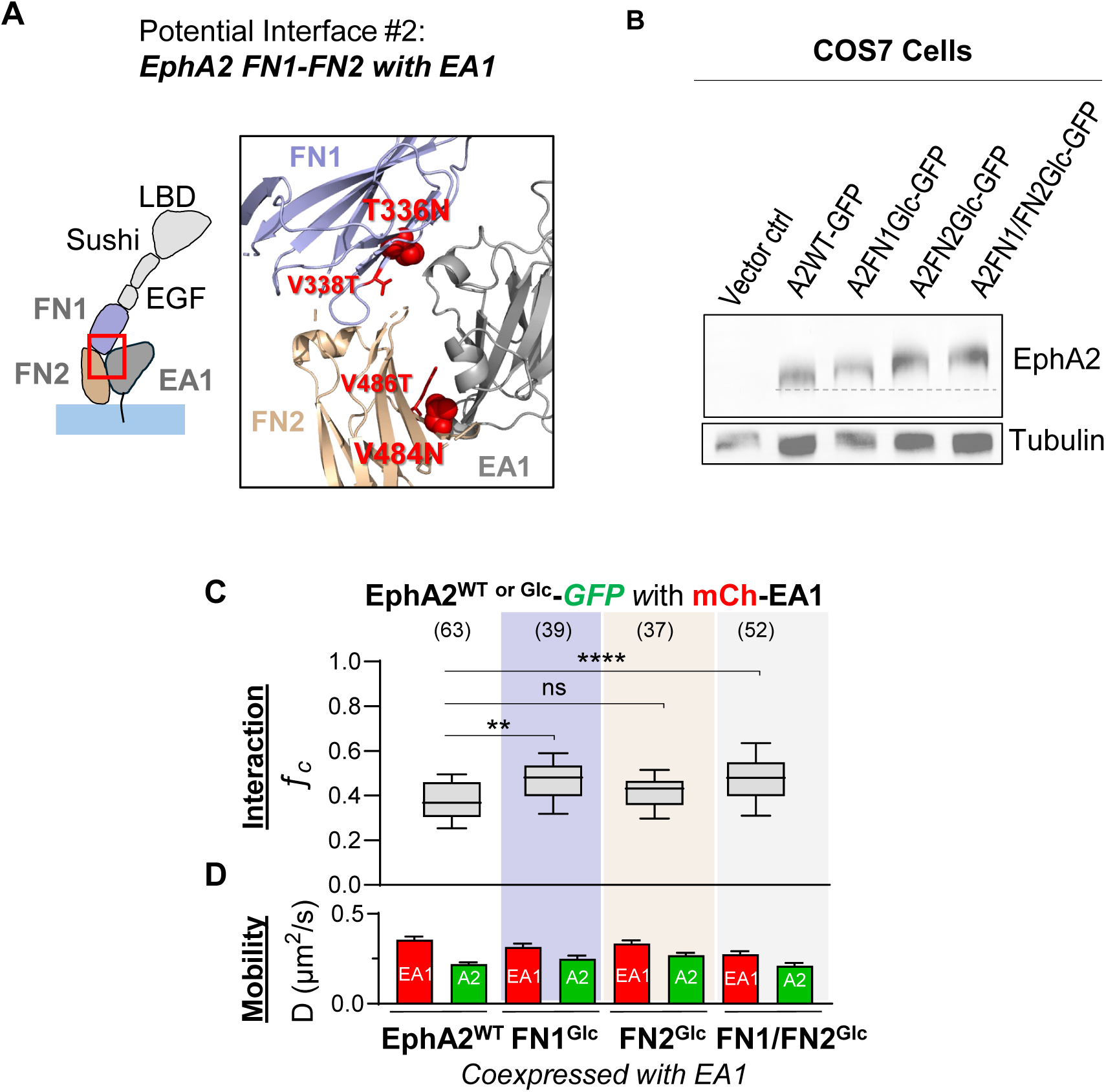
A) Schematic depiction of the potential tripartite interaction between ephrin-A1 and the EphA2 FN1-FN2 domains as observed in the crystal structure. Mutated residues for glycosylation induction are colored in red. Sites for glycosylation are highlighted as red spheres. B) EphA2WT, FN1Glc, FN2Glc, FN1/FN2Glc constructs were transiently expressed in COS7 cells. Whole cell lysates (WCL) were collected and subjected to western blot. C) fc values and D) diffusion coefficients of mCh-EA1 co-expressed with either EphA2WT-GFP (white area) or EphA22Glc-GFP in which one or two glycosylation sites are introduced (purple area: on FN1 domain, beige area: on FN2 domain, grey area: on both FN1 and FN2 domains). EphA22Glc does not abolish the cis EA1-EphA2 interaction and enhanced it instead. The box plots in panel B represent third quartile, median, and first quartile, and the whiskers indicate the 10th to 90th percentile. The total cell number that was used for each sample is reported at the top of the box plots. Data were analyzed by T-test; ****p < 0.0001, **p < 0.001, *p < 0.01.

To verify the successful introduction of glycosylation sites, the N336-, N484-, or N336/N484-EphA2 were *trans*fected into Cos7 cells. Immunoblot of cell lysates showed an upshift upon introduction of each glycosylation site individually (N336 in FN1 or N484 in FN2) compared with EphA2^WT^, and a further upshift when two sites were combined, consistent with successful glycosylation on the respective sites (**Fig. 3B**). Treatment with peptide N-glycosidase (PNGase F) to strip the glycosylation restored the molecular weights of all mutant EphA2 to that of the WT, further confirming successful introduction of glycosylation on both sites (**Fig. S3A**).

Next, we coexpressed ehrin-A1-mCherry with WT-, N336-, N484- and N336/N484-EphA2-GFP, and performed PIE-FCCS measurements. As shown in **Fig. 3C**, introduction of glycosylation on FN1, FN2, or their combination (FN1/FN2) failed to reduce the *f_c_* values or increase D values (**Fig. 3C, D**), suggesting the mutations did not disrupt the heterotypic receptor-ligand interactions *in cis*. On the contrary, an unexpected increase in *f_c_* values was observed (**Fig. 3C**). This likely stems from the disruption of the Head-Tail interface that contributes to the auxiliary arms of EphA2 multimeric assembly as reported previously and illustrated in **Fig. S3B** (16). The bulky glycosylation on FN1 and FN2 may sterically interfere with the auxiliary assembly, freeing up EphA2 monomers to participate in EphA2-Ephrin-A1 interactions *in cis* (**Fig. S3C**). We conclude that the EphA2-ephrin-A1 interaction *in cis* is primarily driven by the LBD and RBD domains, while the FN2 domain and the tripartite interactions do not play a significant role.

### EphA2-ephrin-A1 *cis* interaction competes with *trans* interaction

Having established that EphA2 and ephrin-A1 interactions *in cis* and *in trans* share the same interface, next we investigated if there is a competition between these two modes of interactions. mCh-ephrin-A1^WT^ and EphA2-GFP were coexpressed in cells, and soluble monomeric ephrin-A1 (*trans*EA1) was added to mimic ligand presented *in trans*. Their interaction was examined using PIE-FCCS. Upon addition of soluble *trans*EA1, we observed a significant decrease in *f_c_* value from 0.34 to 0.20 (**Fig. 4A**), indicating weakened interactions between the EphA2 and ephrin-A1 coexpressed on cell surface. Consistently, mCh-ephrin-A1^WT^ showed faster mobility upon *trans*EA1 addition (**Fig. 4B**, red). EphA2-GFP showed slower motility (**Fig. 4B**, green), which was expected as the *trans*EA1 binding to EphA2 is known to induce the formation of large receptor-ligand clusters (1, 13). Together these results suggest a competitive relationship between the *cis* and *trans* interactions (**Fig. 4C**). However, *cis* interaction was not completely abolished, likely due to an equilibrium between the two binding states.

**Figure 4.**
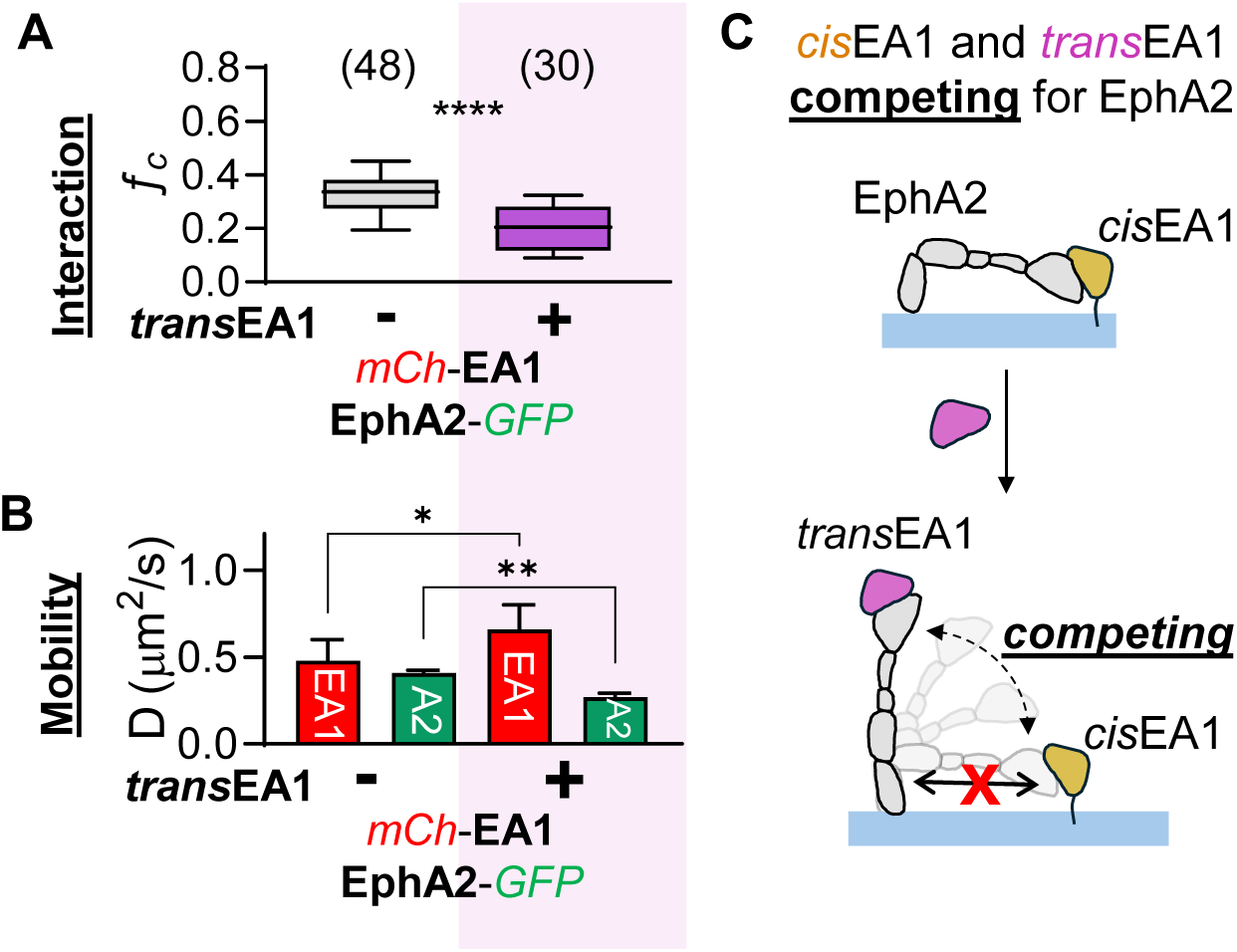
A) fc values and B) diffusion coefficients of mCh-EA1 co-expressed with EphA2-GFP without (white area) and with transEA1 treatment (purple area). Treatment of transEA1 weakens cisEA1-EphA2 interaction. The box plots in panel B represent third quartile, median, and first quartile, and the whiskers indicate the 10th to 90th percentile. The total cell number that was used for each sample is reported at the top of the box plots. Data were analyzed by T-test; ****p < 0.0001, **p < 0.001, *p < 0.01. C) Cartoon depicting the competition between cisEA1 and transEA1 with EphA2.

### EphA2-ephrin-A1 interaction in cis inhibits activation of EphA2 canonical signaling by ligands presented *in trans*

To determine the functional implications of the *cis* interaction, we coexpressed EphA2 with either ephrin-A1^WT^ or ephrin-A1^KE/EK^. For these experiments, we used 283LM cell line derived from a chemically induced squamous cell carcinoma on an *Efna1;Efna3;Efna4* triple knockout (TKO) mouse with genomic deletion of three of the five mammalian *Efna* genes coding ephrin-A ligands (16, 20). These cells also express undetectable levels of ephrin-A2 and ephrin-A5, providing a clean background for the experiments. The 283LM cells co-expressing EphA2 together either mCh-ephrin-A1^WT^ (*cis*EA1WT) or mCh-ephrin-A1^KE/EK^ (*cis*EA1^KE/EK^) were plated at low density to minimize cell contact-induced *trans* interactions.

As shown in **Fig. 5A**, when EphA2 was expressed alone, stimulation with soluble ephrin-A1 mimicking *trans* interaction (*trans-*ephrin-A1) induced dose-dependent activation of canonical signaling, which was marked by tyrosine phosphorylation (pY588) and suppression of Akt phosphorylation (**Fig. 5A**) (12, 16, 20, 21). In contrast, when ephrin-A1WT was coexpressed to establish the *cis* interaction (*cis-*ephrin-A1WT), EphA2 showed diminished activation of canonical signaling and reduced suppression of pAkt. Remarkably, when *cis* interaction was disrupted by the ephrin-A1^KE/EK^ mutation, EphA2 responded strongly to *trans*-ephrin-A1 stimulation, which is comparable with cells expressing EphA2 alone (**Fig. 5A**). These results demonstrate the EphA2-LBD/ephrin-A1 RBD *cis* interaction impedes EphA2 canonical signaling induced by ligands presented *in trans*.

**Figure 5.**
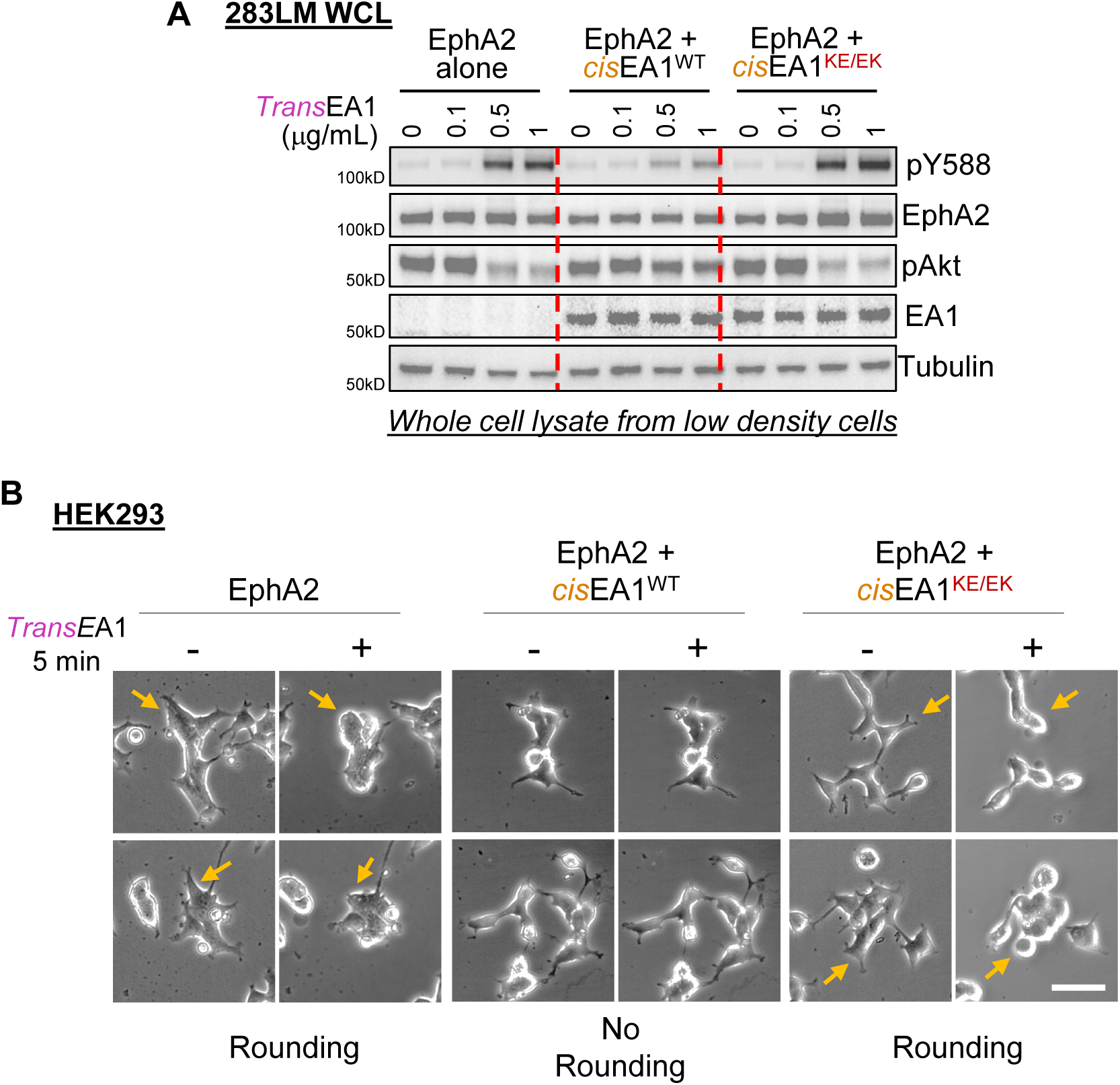
**A)** 283LM cells expressing EphA2 and EA1^WT^ or EA1^KEEK^ were stimulated with different concentrations of ephrinA1 (*trans*EA1) for 15 min and lysed. The whole-cell lysates were subjected to immunoblotting with the indicated antibodies. EA1-EphA2 *cis* interactions reduced responsiveness to EA1 presented in *trans*. **B**) Representative images of HEK293 cells expressing EphA2 and EA1^WT^ or EA1^KE/EK^. Images were taken at 0 min and 5 min after treatment with ephrinA1. Yellow allows indicate cells before and after *trans*EA1 treatment.

A well-documented *in vitro* hallmark of cellular response to EphA2 activation is cell rounding (13), which is in keeping with the extensively documented role of Eph-ephrin interactions in suppression of membrane protrusions and repulsion of migrating cells(13, 22, 23). In cells expressing EphA2 alone, *trans-*ephrin-A1 stimulation triggered rapid cell rounding within 5 minutes (**Fig. 5B**). Consistent with the lack of response to *trans*-ephrin-A1-indcued canonical signaling, cells coexpressing *cis*EA1WT and EphA2 showed no such morphological changes, supporting the inhibitory role of the *cis* interaction. The effects were abolished by KE/EK mutation of ephrin-A1 (**Fig. 5B**). Together, these results demonstrate that the *cis* interaction between EphA2 and ephrin-A1 suppresses *trans* interaction, leading to critical functional consequences.

## Discussion

By combining PIE-FCCS platform to measure membrane protein interactions in live cells, mutagenesis, and biochemical and cellular characterization, we report here strong *cis* interactions between EphA2 and ephrin-A1 through the LBD and RBD domains, respectively. The interaction sequesters EphA2 in an inactive state and renders it significantly less responsive to ligand stimulation *in trans*. EphA2 and ephrin-A1 are the most abundantly coexpressed pair in normal epithelial cells in vitro and in vivo, and are often dysregulated in disease processes, particularly in cancer. The detailed molecular and functional characterization of *cis* interaction could have implications in our understanding of dynamic crosstalk and therapeutic utilities of the pair.

Multiple studies in vitro and in vivo provide functional significance and molecular basis for the Eph-ephrin *cis* interaction. However, significant discrepancies persist as to how the *cis* interactions are assembled. Drescher and colleagues are the first to report that ephrin-A2 and -A5 exert an inhibitory function on their coexpressed receptors EphA3,4,5 in the developing retina (7). Later studies by the same team mapped the interaction interface to FN1-FN2 domains of EphA3 and RBD of ephrin-A5. Using the spinal cord axon guidance model, Kao and Kania also provide evidence that FN2 of EphA4 is involved in interacting with ephrin-A5 and ephrin-B2 (4). In cancer cells, the *cis* interactions between EphA3 and ephrin-A3 are suggested to be mediated by FN1 and FN2 domains (3). Moreover, inspection of the co-crystal structure of EphA2-ephrin-A5 complex also revealed extensive contact surface between ephrin-A5 and FN1 and FN2 domains of EphA2. While these studies using a variety of systems and different receptor/ligand pairs are consistent with the involvement of FN1 and FN2 domains, Yin et al. report that it is LBD of EphA4 that mediates the *cis* interactions with ephrin-A2, which is supported by a function-blocking anti-ephrin-A2 monoclonal antibody that diminishes the *cis* interaction (9). Adding to the complexity, one study suggests that the coexpressed EphA4 and ephrin-A5 are segregated into distinct membrane microdomains and do not engage in apparent *cis* interaction (11). Of note, no attempt has been made to molecularly pin down the interactions between the specific amino acids that are involved in the *cis* interactions.

In the current study, we used PIE-FCCS technology for direct biophysical measurement of protein interactions in live cells and revealed strong EphA2-ephirn-A1 interactions *in cis* through the LBD and RBD domains, respectively. Combining molecular modeling and mutagenesis approaches, we were able to map two salt bridges that partake in the contact interface. Interestingly, the interface is shared with those mediating interactions *in trans* that induces canonical signaling of EphA2. The EphA2^FN1-FN2^-ephrin-A1 interface, that has bene suggested to mediate in the *cis* interactions in several previous studies, does not seem to play a major role, as disruption of the tripartite interface observed in the crystal structure stimulated EphA2-ephrin-A1 *in cis* interaction, rather than reduced it. The reason behind the differences between our results and previous reports remain unclear. One potential explanation is the truncation mutation and co-immunoprecipitation approaches utilized previously. For example, the truncated EphA3 may still bind ephrin-A3 (3) or ephrin-A5 (5) through its co-receptors.

Notably, no previous studies that systematically investigated the *cis* interaction between EphA2 and ephrin-A1. Among the nine Eph receptors and eight ephrin ligands, EphA2 and ephrin-A1 are the most abundantly coexpressed receptor/ligand pair in many epithelial cells in vitro and in vivo (24, 25). Importantly, this interaction has profound functional implications for EphA2, one of the most extensively studied Eph kinases, as it silences EphA2-medicated canonical signaling, thereby promoting non-canonical signaling that plays a critical role in malignant progression. Our previous works elucidated the molecular mechanism for the dual function of EphA2 (16), demonstrating that it can mediate both tumor-suppressive and oncogenic signaling depending on its ligand binding state. Our current data show that *cis* interaction between EphA2 receptors does not trigger activation but instead lead to non-productive interaction and the receptor accumulation on the cell surface. Such non-productive interaction facilitates non-canonical signaling, which has important implications for tumorigenesis and metastasis (16). EphA2 is often overexpressed on cancer cells. Since the affinity of the *cis* interaction is comparable to that of the *trans* interaction due the shared interface, it effectively sequesters ephrin-A1 in the complex, attenuating its ability to engage EphA2 on adjacent cells, further reducing the canonical signaling in the tumor microenvironment and promoting tumor progression.

This study further exemplifies how *cis* and *trans* interactions can lead to distinct functional outcomes; a phenomenon previously observed in multiple immune-related cell surface receptors. For instance, the inhibitory receptor Ly49A on natural killer cells bends over to interact with co-expressed H2-Dd (26). PD-L1 and PD1 (27) or CD28 and CD80 (28, 29) can also be co-expressed and interact on the same cell. These cases illustrate that *cis* interactions can either (1) pre-occupy the receptor, thereby preventing canonical *trans* interactions (27, 29, 30) or (2) induce cell-intrinsic signaling and bypass the sparse opportunity for *trans* interactions and directly stimulates receptor activity (31). However, these earlier studies were often challenged by technical limitations, particularly the difficulty of isolating *cis* interaction from *trans* interaction. However, advanced fluorescence-based techniques such as FCCS (32) and FRET (28, 29, 33), combined with biochemical assays, enabled the validation of *cis* interactions and the significance of their signaling consequences, which can be readily applied to other *cis* vs interactions.

Finally, the suppression of EphA2 canonical signaling via *cis* interaction with ephrin-A1 has important pathological and therapeutic implications. Imbalance of EphA2 and ephrin-A1 expression is a hallmark of many solid tumors. Late-stage malignant tumors frequently exhibit EphA2 overexpression and ephrin-A1 downregulation (2, 34, 35). The sparse opportunity for EphA2-ephrin-A1 *trans* interaction coupled with competition between *cis* and *trans* interaction may dampen the canonical tumor suppressive signaling and contribute to therapeutic resistance. Disrupting the EphA2-ephrin-A1 *cis* interaction while inducing their *trans* interaction can be a promising therapeutic strategy.

## Materials and Methods

### Recombinant proteins

ephrinA1-Fc (Miao et al., NCB, 2000 (ref)); Recombinant human ephrin-A1 protein (His Tag) (SinoBiological, #10882-H08H).

### Antibodies

Rabbit monoclonal Anti-phospho-EphA2 (Tyr588, Cell Signaling Technology, #12677S); Rabbit monoclonal Anti-phospho-EphA2 (Cell Signaling Technology, #6997S); Rabbit polyclonal Anti-ephrin-A1 (Santa Cruz Biotechnology, #sc-377362); Mouse monoclonal Anti-αTubulin (R&D Systems, #MAB9344-100).

### Plasmid

Human EphA2 cDNA (AA 1-971 based on NM_004431.4) with C-terminus GFP was purchased from Origene in gateway donor vector. Human EFNA1 cDNA (AA1-70 based on NM_004428.2) with N-terminal GFP or mCherry tag were purchased from GeneCopoeia in gateway donor vector. The donor vectors were amplified with NEB 5α competent E. coli (New England BioLabs, #C2987). The donor vectors containing EphA2 constructs were shuttled into pLenti and pBabe gateway destination vectors with puromycin or hygromycin selection marker using Gateway LR Clonase II (ThermoFisher Scientific, #56484). The pLenti or pBabe gateway destination vectors were obtained from Addgene (#17452, #17454, #51070). The resulted pLenti or pBabe vectors containing EphA2 constructs were amplified with NEB stable competent E. Coli (New England BioLabs, #C3040) and were used for Lenti/Retro virus production.

### Cell lines

Human HEK-293(ATCC, CRL-1573); Cercopithecus aethiops COS-7 (ATCC, CRL-1651); Mouse 283LM (16); Human Pheonix-AMPHO (ATCC, CRL-3213).

### Cell Transfection

Plasmids with pLenti vectors were *trans*fected into COS7 cells using lipofectamine 2000 (Invitrogen #11668027). The resulted COS7 cells expressing EphA2 and/or ephrinA1 WT/mutants were used in PIE-FCCS measurements.

### Retro-virus Mediated Gene Transduction

Plasmids with pBabe vectors were *trans*fected into Phoenix retroviral packaging cells with lipofectamine 2000. HEK293, 283LM cells were infected with retroviral mediated gene *trans*fer in the presence of 6 mg/mL polybrene, selected in the presence of puromycin, and sorted with SONY MA900 FACS sorter before analysis.

### Ligand Stimulation, Whole Cell Lysate Collection, and Western Blot Analysis

Monodispersed 100,000 HEK293 or 283LM cells were plated on 10-cm cell culture dish and incubated overnight. The cells were stimulated with 0-1.0 ug/mL monomeric ephrin-A1. At 15 minute, cells were lysed in modified RIPA buffer (20 mM Tris, pH 7.4, 120 mM NaCl, 1% Triton X-100, 0.5% sodium deoxycholate, 0.1% SDS, 10% glycerol, 5 mM EDTA, 50 mM NaF, 0.5 mM Na_3_VO_4_, phosphatase inhibitor cocktail, and protease inhibitor, including 1 mM phenylmethylsulphonyl fluoride, 2 mg/mL of aprotinin, and 2 mg/mL of leupeptin). Lysates were incubated for 30 min on ice and then centrifuged at 13,000g for 10 min at 4 °C, and either analyzed immediately or stored at -80 °C. Whole cell lysates were separated on Bolt 4-12% Bis-Tris Plus gel (Invitrogen, #NW04125BOX) and electro*trans*ferred onto polyvinylidene difluoride membranes (Milipore, #IPVH00010), which were then blotted using indicated antibodies. The same membrane was used for blotting different antibodies after stripping the previous blot.

### Glycosylation verification

COS7 cells were plated on 6 well plate and incubated overnight. GFP-tagged EphA2 WT, FN1Glc, FN2Glc, FN1FN2Glc plasmids with pLenti vectors were *trans*fected into the cells using lipofectamine 2000. 48 hours later, GFP expression was verified under the fluorescence microscope, and cells were lysed using cell lysis buffer (Cell Signaling Technology, #9803) with phosphatase inhibitor cocktail and 1 mM phenylmethylsulphonyl fluoride. EphA2 WT or mutant proteins were immunoprecipitated using ephrinA1-Fc and Protein G Magnetic Beads (Cell Signaling Technology, #70024) according to the manufacturer’s manual. To remove glycosylation, beads were incubated with Peptide N-Glycosidase F (New England Biolabs, #P0704S) according to the manufacturer’s manual. The original or deglycosylated proteins were subjected to immunoblotting for confirmation.

### Live-cell Imaging

HEK293 cells were plated on 24-well cell culture dish. During cell rounding assay, the culture dish was kept in INU WSKM on-stage incubation chamber (Tokat Hit) at 37 oC and 5% CO2. 0.5 μg/ml ephrin-A1-Fc were added to stimulate cells. Bright field images were taken on Leica DMi8 Microscope.

### Pulsed Interleaved Excitation - Fluorescence Cross-Correlation Spectroscopy (PIE-FCCS) Instrumentation and data analysis

PIE-FCCS measurement was performed on a customized Leica DMi8 inverted microscope (Leica Microsystems Inc., Deerfield, IL) with home-built pulsed interleaved excitation and time-correlated single-photon dection. A 5-picosecond-pulsed supercontinuum white light laser (SuperK Flanium FIR-20, NKT Photonics Inc., Boston, MA) was used as excitation laser source. The custom-built excitation box for pulsed interleaved excitation and custom-built emission box for time - correlated single photon detection was set as previously described (16). Data recorded by TCSPC card is input to a computer for correlation with home-written Matlab scripts. For live cell PIE-FCCS experiments, cells were plated onto MatTek glass bottom culture dish 24-48 hrs before the experiments. Cell culture media was displaced by pre-warmed Opti-MEM before the PIE-FCCS measurement, and the temperature was maintained at 37 oC via on-stage incubator throughout the measurement. For stimulation, 1 ug/mL ligand was added and incubated for 5 min prior to measurement. To analyze the data, time-tagged photon counts were split into 10-second segments, then processed through binning and gating to eliminate spectral crosstalk. A custom MATLAB script was used to compute auto- and cross-correlation curves for each molecular species. The resulting correlation functions from each measurement were averaged and fitted to a previously described 2D single-component diffusion model. Average Dwell time (τD) from the curve fit was used to calculate the effective diffusion coefficient (D=ωo2/4τD). Using the ratio of auto- to cross-correlation amplitudes, a cross-correlation value (ƒc) was determined for each cell to evaluate the receptor-ligand interaction. Detailed description of the calculation is described previously in detail (16).

## Acknowledgments

We thank MetroHealth Foundation and Dr. Adam W. Smith for support of the instrumentation. This work was funded by NCI training grant T32CA059366 (X.S.); National Institutes of Health (NIH) grants R01NS096956, R01CA250067, R01CA297778 (BW).

## Author Contributions

SK, XS, BW designed the research and wrote the manuscript. ES provided structural insight and designed the structure-guided mutagenesis strategies, SK, XS, PT and WW performed the research.

