## Supplementgal Figures for "EphA2 and Ephrin-A1 Utilize the Same Interface for Both *in cis* and *in trans* Interactions That Differentially Regulate Cell Signaling and Function"

**Fig. S1**

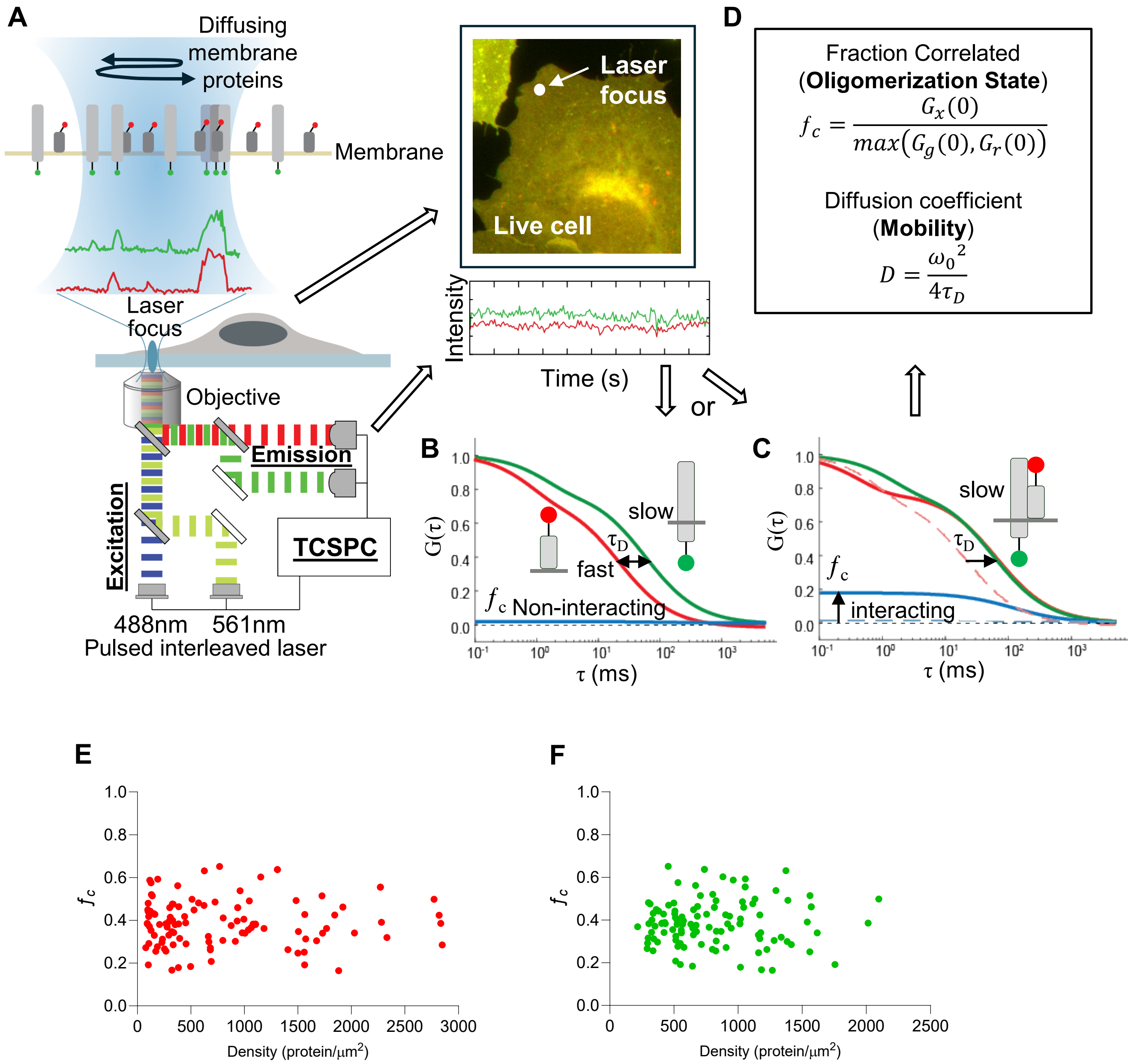


**Figure S1. A**) Schematics of PIE-FCCS instrumentation. Two pulsed interleaved lasers are overlapped and focused on the peripheral membrane of COS7 cells expressing GFP-/mCh-tagged protein of interest (epifluorescence image). The fluorescence fluctuation is recorded and are subjected to auto-/cross-correlation function. **B-C**) Autocorrelation (green and red) curves and cross-correlation curve are obtained. **D**) Two main parameters are calculated from the curves: Cross-correlation value or fraction correlated ($f_c$) and diffusion coefficient (D). **E**) Cross-correlation values ($f_c$) of mCh-ephrin-A1 plotted against molecular density. **F**) Cross-correlation values ($f_c$) of EphA2-GFP plotted against molecular density. Both **E** and **F** shows that there was no density-driven interaction within this expression range.

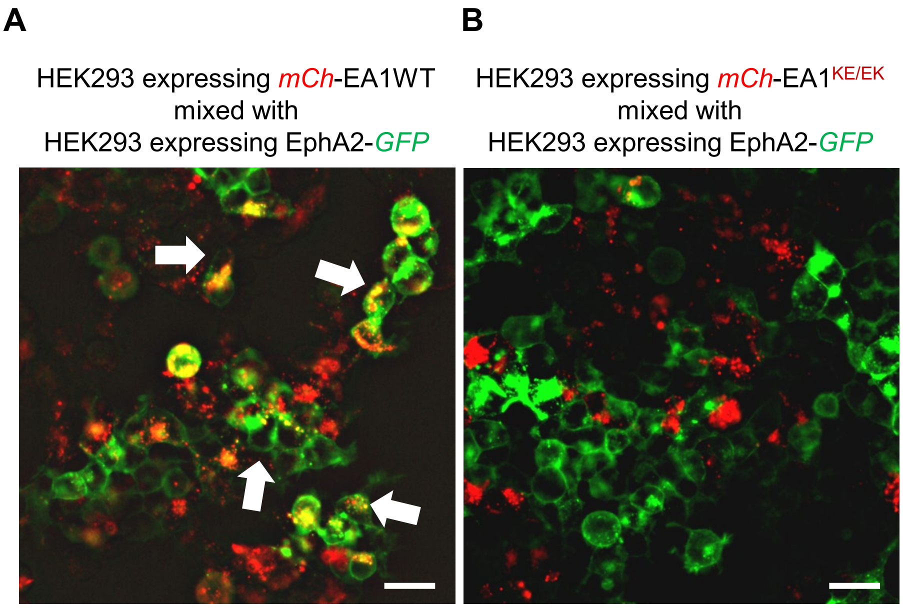


**Figure S2.** A) Fluorescence images of mix of mCh-EA1WT-expressing HEK293 cells and EphA2-GFP-expressing HEK293 cells. B) Fluorescence images of mix of mCh-EA1KE/EK-expressing HEK293 cells and EphA2-GFP-expressing HEK293 cells. These images show that $EA1^{KE/EK}$, which harbors mutations to disable the binding with LBD, cannot activate EphA2. White arrows in the left image point to puncta (orange), which is the indication of ephrin-A1-induced EphA2 activation and endocytosis, whereas in the right image, there is no visible puncta in the EphA2-expressing cells. Calibration bar: 10 μm.

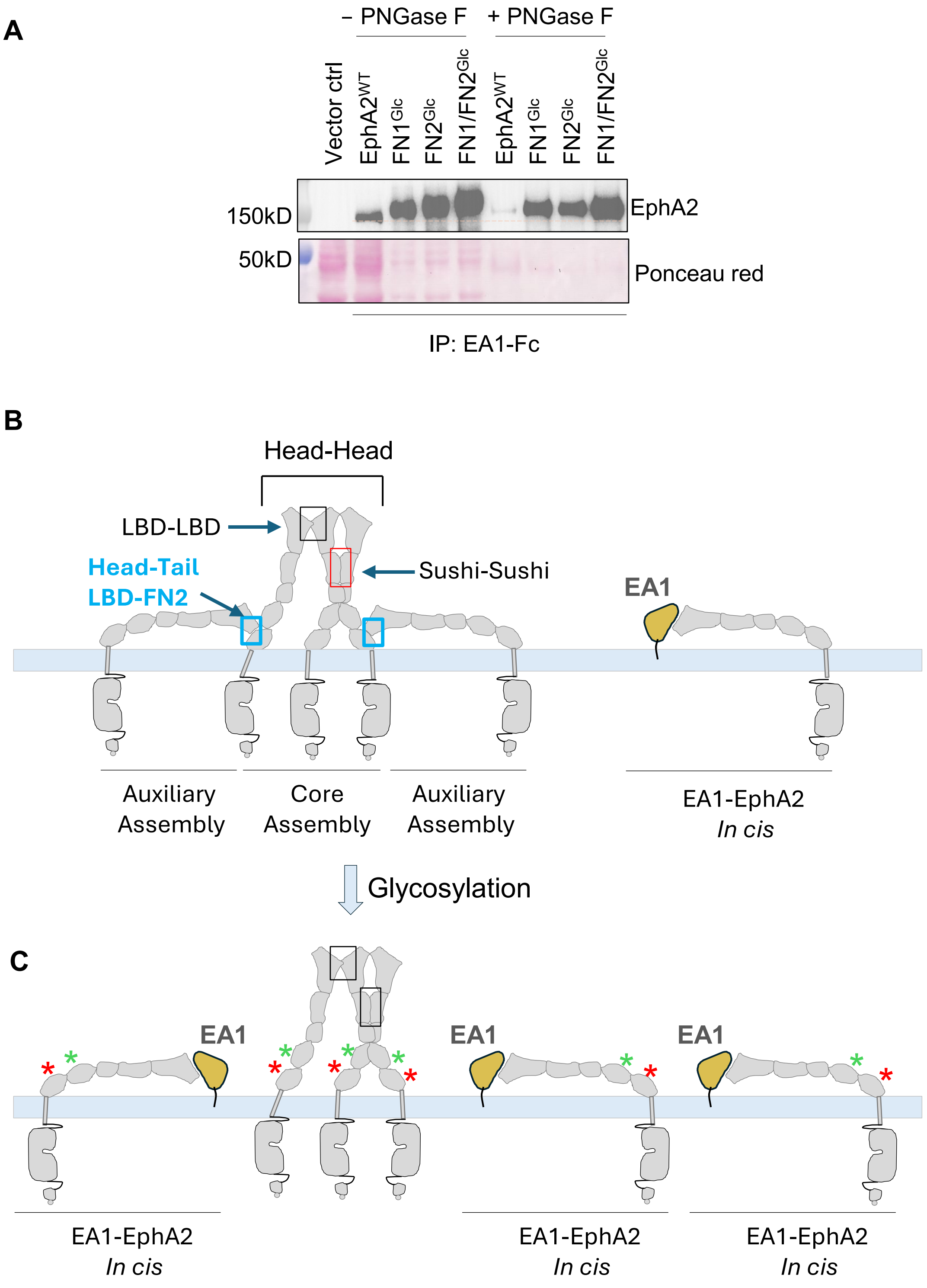


**Figure S3. A)** EphA2$^{WT}$, FN1$^{Glc}$, FN2$^{Glc}$, FN1/FN2$^{Glc}$ constructs were transiently expressed in COS7 cells. Whole cell lysates (WCL) were collected, and EphA2 from each WCL was immunoprecipitated (IP) using EphrinA1-Fc (EA1-Fc), followed by treatment with PNGase F for deglycosylation. The PNGase F-treated and untreated IP products were subjected to immunoblotting to confirm the introduction of glycosylation after mutation. **B-C)** Schematic illustration showing how introduction of glycosylation could disrupt the Head-Tail interaction in the EphA2 multimeric assembly of EphA2. **B)** Multimeric assembly of EphA2 apo receptor as reported recently[16]. **C)** Steric interference by glycosylation disrupts the auxiliary assembly of EphA2, freeing up more EphA2 to engage in interaction with ephrin-A1 *in cis*.
